# RIPK3 promoter hypermethylation in hepatocytes protects from bile acid induced inflammation and necroptosis

**DOI:** 10.1101/2021.01.15.426790

**Authors:** Jessica Hoff, Ling Xiong, Tobias Kammann, Sophie Neugebauer, Julia M. Micheel, Mohamed Ghait, Sachin Deshmukh, Nikolaus Gaßler, Michael Bauer, Adrian T. Press

**Author notes:** **Correspondence** Adrian Press, Am Klinikum 1, 07747 Jena, +49 3641/ 9 323139.

## Abstract

**Background & Aims:** Necroptosis facilitates cell death in a controlled manner and is employed by many cell types following injury. It plays a major role in various liver diseases, albeit the cell type-specific regulation of necroptosis in the liver and especially hepatocytes has not yet been conceptualized.

**Approaches & Results:** Here, we demonstrate that DNA methylation suppresses RIPK3 expression in human hepatocytes and HepG2 cells. In diseases leading to cholestasis the RIPK3 expression is induced in mice and humans in a cell-type specific manner. Over-expression of RIPK3 in HepG2 cells leads immediately to RIPK3 activation by phosphorylation that is further modulated by different bile acids.

**Conclusion:** Bile acids mediated RIPK3 activation facilitates the secretion and expression of IL-8 via the JNK-pathway, suggesting hepatocytes suppress RIPK3 expression to protect themselves from bile acid induced necroptosis and inflammation but in chronical liver diseases associated with cholestasis induction of RIPK3 expression may be an early event signaling danger and repair through release of IL-8.

**Graphical abstract:** 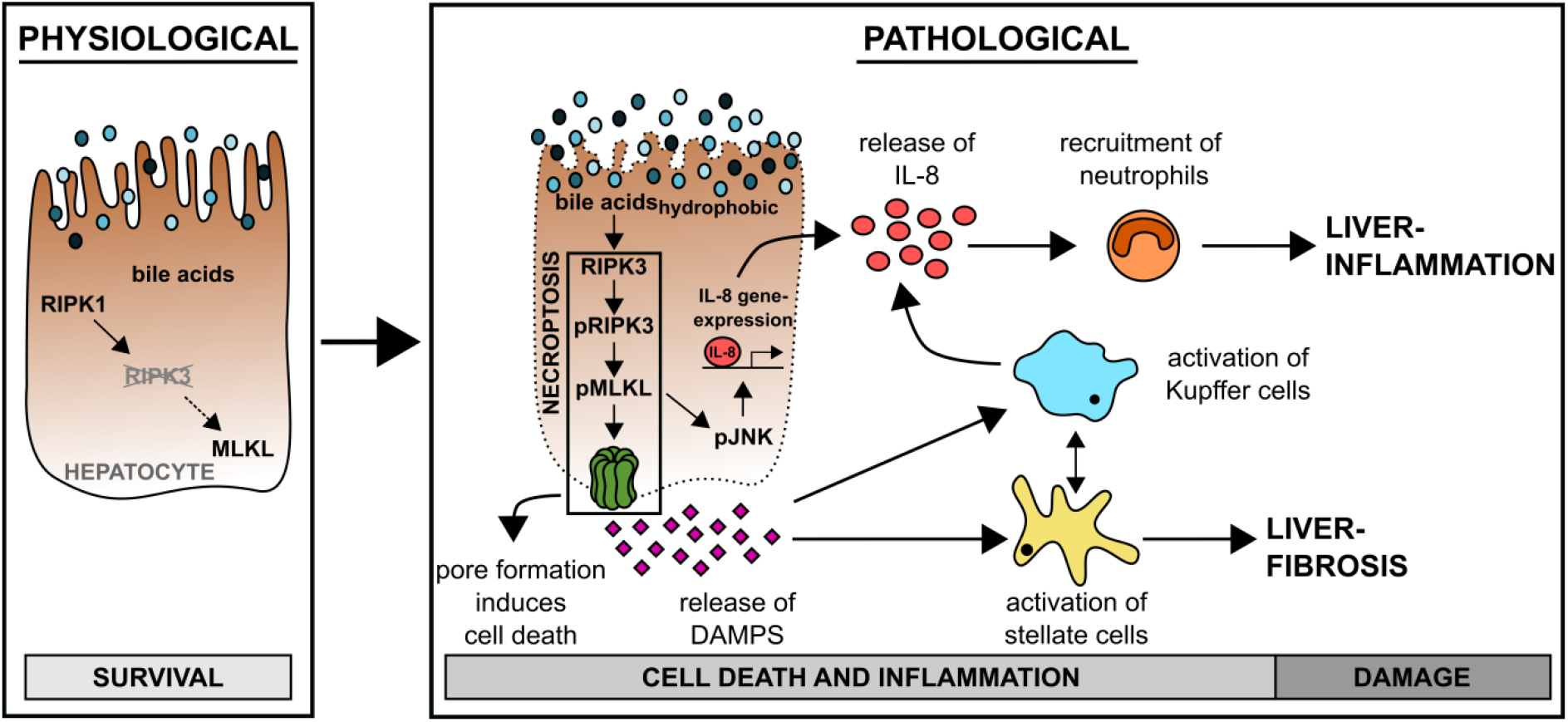

## Introduction

Regulated cell death signaling events, e.g. apoptosis, necrosis, ferroptosis, necroptosis or pyroptosis, are crucial events for maintenance of tissue homeostasis. These pathways are involved in the induction and modulation of the immune response [1, 2] and tissue regeneration [3, 4], where each component contains their specific regulatory mechanism and molecular components. [5]

Necroptosis is a pro-inflammatory cell death type that signals damage and repair to many cell types following injury. [6, 7] Mechanistically, formation of the necroptosis-defining necrosome requires phosphorylation of receptor-interacting serine/threonine-protein kinase 3 (RIPK3) [8] and activation of mixed lineage kinase domain-like protein (MLKL) leading to pore formation and cell death. [9-11] The first steps in the induction of necroptosis are shared with the apoptotic pathway. It has been reported that receptor-interacting serine/threonine-protein kinase 1 (RIPK1) is recruited to an active cytokine receptor, such as tumor necrosis factor receptor type-1 (TNFR1) where it is phosphorylated (pRIPK1) in a multiprotein complex, also known as complex I. [12, 13] pRIPK1 interacts with Fas-associated death domain protein to activate caspase 8 (CASP8) inducing apoptosis. In the presence of CASP8 inhibition, RIPK3 is phosphorylated (pRIPK3) and this represents the first step of the necroptosis pathway. [14] pRIPK3 then associates in a multi-protein complex with MLKL, resulting in its phosphorylation (pMLKL). The pRIPK3/pMLKL complex is then recruited to a membrane where multiple pMLKL proteins oligomerize into a pore, ultimately leading to cell lysis. [15]

Necroptosis is a promising target for future tumor therapeutics [16], however its role in the liver is controversial. [17-22] Studies have cast doubt on hepatocyte RIPK3 expression under physiological conditions. [23] By contrast, hepatocyte injury was reduced in RIPK3 knockouts in varied murine models of acute and chronic liver injury (ethanol-induced [18], ischemia-reperfusion [20], non-alcoholic fatty liver disease [22], and concanavalin-A hepatitis [24]), albeit no protection was seen against injury induced by acetaminophen. [25, 26] Also, necroptosis is emerging as a critical mechanism in pathogenesis of cholestasis. [27] In cholestatic liver disease bile acids are retained and the bile flow is disrupted. [27]

Bile acids are very abundant metabolites in hepatocytes that are known to be involved in the development of different liver diseases. [28, 29] Altogether 15 bile acids are detected in humans and their formation is the primary pathway of cholesterol catabolism which is tightly regulated within the liver parenchyma to prevent the cytotoxic accumulation of bile acids. [30, 31] Cholic acid (CA) and chenodeoxycholic acid (CDCA) are primary bile acids, described as the dominant but not exclusive forms present in the liver. Hepatocytes conjugate CA and CDCA to taurine or glycine during biotransformation before they are secreted into the bile. [31] In the colon bile acids are subjected to various microbial-mediated transformations including deconjugation and transformation of primary to secondary bile acids (lithocholic acid (LCA), ursodeoxycholic acid (UDCA)). [32] Owing to their structural formation, bile acids are classified by their hydrophobicity (hydrophobic: LCA > CDCA > CA > UDCA). [33] The hydrophobic bile acids (LCA, CDCA) are commonly known as toxic bile acids and potent inducers of apoptotic or necrotic cell death, whereas the hydrophilic bile acids are often described as cytoprotective. [34-36]

Besides the well-known function of bile acids as detergents in the digestive tract and signaling under physiologic conditions, they are also highly active signaling molecules for eukaryotic cells in supraphysiological concentrations as they occur in various liver diseases. The importance of regulating inflammation has been highlighted, for instance, in their ability to trigger inflammation and cell death. [37, 38] Presence of pathological concentrations of bile acids in hepatocytes, by accumulation, induces different cell death mechanisms (e.g. apoptosis, necrosis or necroptosis). [36] This implies that hepatocytes must have acquired an endogenous mechanism to counteract bile acids’ pro-inflammatory and cell-toxic properties, e.g. due to the loss of key mediators of inflammation and cell death.

Here, we investigate the expression profile of hepatocellular RIPK3 protein, responsible for the induction of necroptosis, under physiological and pathological liver conditions. Further, we investigate RIPK3 regulatory mechanisms using bile acids in vitro that may prevent and trigger hepatocellular inflammation and tissue regeneration.

## Materials and Methods

### Cell isolation and culture

HepG2 cells were cultured in Dulbecco’s Modified Eagle Medium containing F12 nutrient mix (DMEM:F12; Biozym) supplemented with 10% fetal calf serum (FCS) and 100 iU penicillin and 100 iU streptomycin at 37°C in a humidified atmosphere of 5% CO_2_. Prior to the day of the experiment, cells were washed with phosphate-buffered saline without calcium and magnesium (PBS) and cultured into either 6-, 12- or 96-well tissue culture plates depending on the experimental conditions.

Plateable, cryopreserved primary human hepatocytes (pHep) from one male donor (18 years, BMI 28.7) were purchased from Lonza, Switzerland. Cells were thawed and cultured according to the manufacturer’s instructions using the recommended hepatocyte culture media provided by Lonza. Hepatocytes were seeded at a density of 2×10^6^ cells per well in a 6-well plate coated with collagen type I (Corning) at 10 µg cm^-2^.

Primary human macrophages (pMФ) were isolated from the whole blood of healthy volunteers with Biocoll separating solution (Merck) according to the manufacturer’s protocol and seeded at a density of 2×10^6^ cells per well in a 6-well plate. After 5 days of differentiation by cultivating in Dulbecco’s Modified Eagle Medium (DMEM; Lonza) supplemented with 10% FCS, 10 ng mL^-1^ M-CSF (ReproTech) and 10 µg mL^-1^ ciprofloxacin (Fresenius Kabi), cells were washed with PBS and subsequently used. Purification of the primary cells was characterized by specific markers (**Fig. 1A**).

**Fig. 1:**
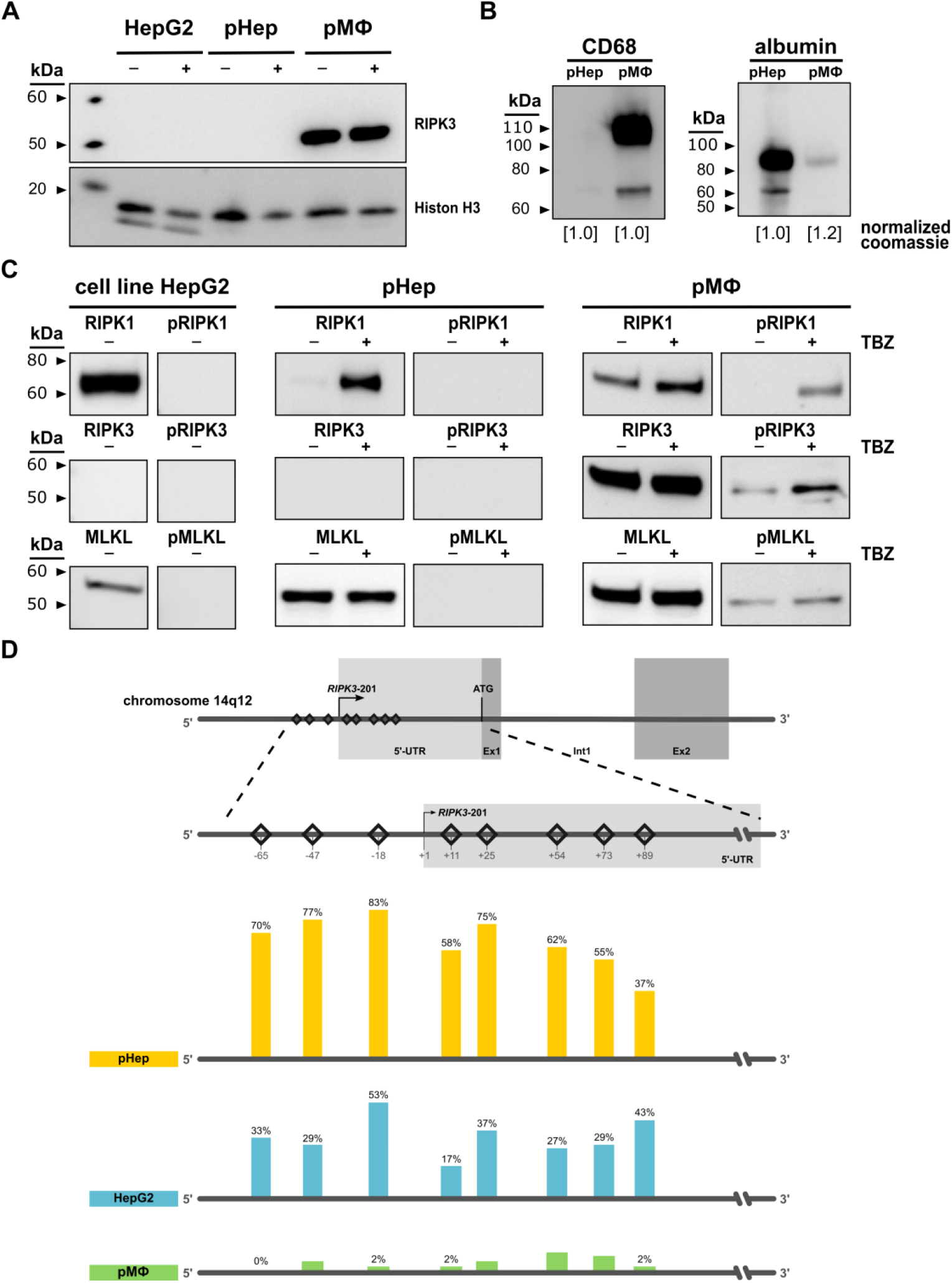
Necroptosis signaling is absent in primary human hepatocytes under physiological conditions. (**A**) Protein expression of RIPK3 in HepG2 cells, primary human hepatocytes (pHep) and primary human macrophages (pMФ). (+) indicates TBZ stimulation with necroptosis inducing reagents where cells were pre-incubated for 0.5 h with BV6 (5 µmol L^-1^) and zVAD-fmk (20 µmol L^-1^) before stimulation with TNF-α (20 ng mL^-1^) for 6 h. Histone H3 and coomassie stained gels were used to control the loading. (**B**) Protein expression of molecules with high abundance in hepatocytes (albumin) or macrophages (CD68) are detected to assess the purity of the primary cells. (**C**) Protein expression of molecules in the necroptosis signaling pathway detected in HepG2 cells, pHep and pMФ. (+) indicates TBZ stimulation with necroptosis inducing reagents where cells were pre-incubated for 0.5 h with BV6 (5 µmol L^-1^) and zVAD-fmk (20 µmol L^-1^) before stimulation with TNF-α (20 ng mL^-1^) for 6 h. n=1-3. (**D**) RIPK3 promoter methylation analysis in pMФ and pHep as well as the hepatocellular carcinoma cell line (HepG2) detected by pyrosequencing. Bars and numbers depict the mean percentage of methylated cytosine in the different promoter regions of each sample. Sequencing was performed on 1 (pHep, pMФ) or 2 (HepG2 cells) individual samples each containing DNA from 1 million cells.

### Transfection

HepG2 cells were transfected with purified plasmid DNA of pcDNA-FLAG or pcDNA3-FLAG-RIPK3 using Lipofectamine 3000 (ThermoFisher Scientific) as a transfection reagent in serum-reduced Opti-MEM medium (ThermoFisher Scientific) according to the manufacturer’s protocol. The pcDNA-FLAG plasmid was purchased from Thermo Fisher (ThermoFisher Scientific). pcDNA3-FLAG-RIPK3 (Addgene plasmid, http://n2t.net/addgene:78815; RRID: Addgene_78815) was a gift from Jaewhan Song. [39]

### Stimulation with Necroptosis-Mix

Cells were washed with PBS and pre-treated with 5 µmol L^-1^ inhibitor of apoptosis protein antagonist BV6 (APExBIO) and 20 µmol L^-1^ pan-caspase-inhibitor zVAD-fmk (Bachem) for 30 min prior to stimulation with 20 ng mL^-1^ tumor necrosis factor-α (TNF-α) (Prospec Inc) in cell culture media.

### Stimulation with Bile Acids

After transfection with pcDNA-FLAG and pcDNA3FLAG-RIPK3 for 24 h, HepG2 cells were carefully washed with PBS and stimulated with DMSO-diluted bile acids ([G/T]CA, [G/T]CDCA, [G/T]UDCA, TLCA: 50 µmol L^-1^; LCA: 5 µmol L^-1^; GLCA: 10 µmol L^-1^) in DMEM:F12 medium (without supplements).

### Mice

Male and female FVB/N and FVB/NRj mice at 8-12 weeks of age were used for all experiments. FVB/N and FVB/NRj mice were partially bred within the animal facility of the Jena University Hospital under specific pathogen-free conditions and purchased from Janvier Labs. The animals had access to conventional rodent chow and water ad libitum. They were maintained under a constant humidity (50-60%), 12-hour light/dark cycle (incl. 20 min dim phases) and a constant temperature (24°C). All experimental protocols were approved by the ethics committee and local government authorities in Thuringia, Germany.

### Experimental mouse models

#### Surgical animal models

Surgery was performed under anesthesia inhalation (1-2% Isoflurane, CP-Pharma and 100 mL min^-1^ carbogen) and 1 mg kg^-1^ body weight p.o. Meloxicam (0.5 mg mL^-1^ suspension, CP-Pharma) for pain-relief was given 1 h before surgery. A midline incision was used to open the abdomen and expose the bile duct and liver. After the surgical procedure (details for different models are given below) the abdominal layers were closed with 4-0 antibacterial suture (Ethicon) and Bupivacaine (2-4 mg kg^-1^ body weight, PUREN Pharma) was administered intra-incisional for topical anesthesia. The animals received Ringer acetate (20 mL kg^-1^, Berlin Chemie AG) subcutaneously for fluid resuscitation and were offered an additional heat source (warm lamp) during the recovery phase. Normal and soaked food was available on the ground for the animals at all times after surgery. The animals were scored and weighed a minimum of twice daily. The scores were designed to reflect post-surgical conditions as well as specific symptoms of surgical intervention.

##### General surgery

This group was used as a sham group. After exposure of the liver and bile duct, the abdominal layers were closed without additional intervention.

##### Bile duct ligation (BDL)

Intrasurgical the bile duct was ligated twice with two 6-0 braided silk sutures (Teleflex Medical).

##### Warm Ischemia Reperfusion injury (IR)

Intrasurgical a microvascular clamp was placed above the left lateral branch of the portal vein to interrupt the blood flow to the left lateral lobe. The liver was covered and kept moist with Ringer acetate while the body temperature was maintained at 37°C with a heating plate. After 60 min of partially hepatic ischemia, the clamp was removed to initiate the reperfusion.

#### Non-surgical animal models

##### Acetaminophen induced liver injury (APAP)

Animals had been fastened for 16 h prior APAP injection to reduce metabolic activity and glutathione levels in the liver. Acetaminophen (Sigma Aldrich) was dissolved in pre-warmed PBS and mice were treated with 300 mg kg^-1^ body weight intraperitoneally. [40] After the APAP injection, mice had free access to food and water. Sham animals received an intraperitoneal injection of PBS.

##### Peritoneal Contamination and Infection (PCI)

Human feces (60-70 µL feces suspension) diluted in Ringer acetate (Berlin Chemie AG) (PCI group) or Ringer acetate alone (sham group) was injected intraperitoneally. The bacterial composition of the human feces suspension, mainly *Escherichia coli* (2.5×10^8^ CFU per mL), *Actinobacteria* (2.5×10^8^ CFU per mL) and *Bifidobacteria* (2.9×10^8^ CFU per mL) used in this study had been characterized previously. [41] Additionally, 2.5 mg of metamizole (Novaminsulfon-Ratiopharm, Ratiopharm) was administered every 6 h on the tongue of the animal. Twice a day animals received subcutaneously 25 mg kg^-1^ body weight meropeneme (Inresa Pharmaceutical, 2.5 mg mL^-1^ diluted in Ringer acetate).

### Hepatocyte isolation

Animals subjected to surgical or non-surgical models of liver injury were sacrificed after defined time points by an overdose of ketamine (> 300 mg kg^-1^ body weight, CP-Pharma) and xylazine (> 50 mg kg^-1^ body weight, Bayer). For hepatocyte isolation a liver perfusion was performed. The system is composed of a peristaltic pump with adjustable speed, a silicone tubing immersed in different buffers in the water bath (38°C) and a cannula (we used 25 G) at the tube outlet. The pump speed was set to a maximum of 5 mL min^-1^ (beginning) and was increased to 12 mL min^-1^ for perfusion after the cannulation of the portal vein. The perfusion was started using Krebs Henseleit Buffer (KHB, Biochrom) supplemented with 8 U mL^-1^ heparin (Heparin-Natrium-2500-ratiopharm, stock solution: 5000 I.E. mL^-1^, Ratiopharm). When the liver appeared pale, the perfusion medium was changed to Liver Digest Medium (LDM, ThermoFisher Scientific). Perfusion was maintained until the liver appeared digested. Livers were separated and kept in a cell culture dish with some LDM after perfusion and incubated for an additional 5 min at 37°C. The tissue was strained through a 70 µm nylon cell strainer (Corning) into a conical tube using approximately 30 mL of DMEM (ThermoFisher Scientific). Hepatocytes were isolated and purified by three times centrifugation for 4 min at 40 g and 4°C. Between each centrifugation step the supernatant was removed and replaced with 30-40 mL DMEM to wash the hepatocyte pellet. Aliquots and lysates were prepared after counting hepatocytes in 10 mL DMEM.

### Methylation Analysis

Genomic DNA was extracted from cell cultures using the DNeasy blood & tissue kit (Qiagen) according to the manufacturer’s protocol. Cells were then re-suspended in the proteinase K digestion buffer (40 µg proteinase K) and incubated at 50°C for 30 min. Cell debris was pelleted by centrifugation at 14000 g for 10 minutes. For DNA methylation analysis, 500 ng of the purified genomic DNA was bisulfite converted using the EZ DNA Methylation-Direct kit (Zymo Research). Bisulfite-treated DNA was purified according to the manufacturer’s protocol and was eluted to a final volume of 46 µL. PCRs were performed using 1 µL bisulfite-treated DNA and 10 µmol L^-1^ of each target-validated primer from the EpigenDx Assay (ADS1678FP, ADS1678RPB). One primer was biotin-labeled and HPLC-purified to purify the final PCR product using sepharose beads. PCR products were bound to Streptavidin Sepharose HP (GE Healthcare Life Sciences) for immobilization. Afterwards the immobilized PCR products were purified, washed, denatured with 0.2 μmol L^-1^ NaOH solution, and washed using the Pyrosequencing Vacuum Prep Tool (Qiagen) as stated in the manufacturer’s protocol. A methylation assay ADS1678 (EpigenDx) was performed, which reports the methylation of eight representative CpG sites in the regulatory region (**Tab. 1**). 10 µL of the PCR products were sequenced by pyrosequencing on the PSQ96 HS system (Qiagen) following the manufacturer’s instructions. The methylation status of each CpG site was determined individually using QCpG software (Qiagen). The software calculates the level at each CpG site as the percentage of the methylated alleles divided by the sum of all methylated and unmethylated alleles.

### Western Blot

After stimulation, cells were washed with PBS prior to lysis in RIPA buffer (150 mmol L^-1^ NaCl, 1 mmol L^-1^ EDTA, 0.1% SDS, 1% Triton X-100, 500 mmol L^-1^ Tris-HCl, 0.5% deoxycholic acid) containing inhibitors of phosphatases (1:10 PhosphoStop; Roche) and proteases (1:100 Protease Inhibitor Cocktail; ThermoFisher Scientific). After determination of the protein concentration by BCA Protein Assay Macro Kit (Serva Gelelectrophoresis GmbH) 10 µg of cell extracts were loaded and separated by SDS-PAGE, transferred to 0.2 µm PVDF membranes (Carl Roth), blocked with 5% bovine serum albumin (BSA) or nonfat dry milk in TBS-T and incubated with monoclonal antibody rabbit IgG (in 5% BSA/ nonfat dry milk in TBS-T) overnight at 4°C. All antibodies were purchased from Cell Signaling Technology targeting either RIPK1 (#3493, 1:750), pRIPK1 (#65746, 1:750), RIPK3 (#13526, 1:750), pRIPK3 (#93654, 1:750), RIPK3 (#95702, 1:750), MLKL (#14993, 1:750), pMLKL (#91689, 1:750), Histone H3 (#4499, 1:2000), JNK (#9258, 1:1000), pJNK (#4668, 1:1000), albumin (#4929, 1:1000), CD68 (#86985, 1:1000) or MCAM (#68706, 1:1000). Membranes were washed with TBS-T and incubated with HRP-conjugated goat anti-rabbit IgG antibody (1:2000 in 5% nonfat dry milk in TBS-T; Jackson ImmunoResearch) at room temperature for 1 h. For visualization, the chemiluminescence method was performed with normal and ultra-sensitivity substrate (SERVA*Light* Eos CL HRP WB Substrate Kit; SERVA*Light* EosUltra CL HRP WB Substrate Kit; Serva Gelelectrophoresis GmbH).

### RIPK3 staining of Human Samples

To investigate the relationship between RIPK3 expression and cholestasis, patients suffering various liver diseases (**Tab. S3**) were classified according to the bilirubin plasma concentration in two groups, no cholestasis (< 21 µmol L^-1^) and cholestasis (≥ 21 µmol L^-1^). Paraffin-embedded liver tissue sections (4 µm thickness) mounted on glass slides were used. For immunostaining, tissue sections were deparaffinized and permeabilized by citrate buffer (pH 6) for 25 min (Dako). Slices were blocked with 5% donkey serum (37°C, 30 min) and then incubated with a primary antibody targeting RIPK3 (#95702, Cell Signaling Technology) at 4°C overnight diluted in antibody diluent (Dako). Negative controls were incubated with antibody diluent only. Slides were then incubated with Alexa-Fluor 568-conjugated anti-rabbit IgG (1:500, ThermoFisher Scientific) and 5 U mL^-1^ DY-636 conjugated Phalloidin (Dyomics) for 60 min at room temperature. Finally, the sections were mounted with Roti-Mount FluorCare mounting media (Carl Roth).

Images were acquired on a laser scanning microscope (LSM-780, Carl Zeiss, Germany) at 400x magnification (40x, numeric aperture 0.95, Carl Zeiss, Germany) and 4.82 pixel per micron. Alexa Fluor 568 fluorescence was excited at 561 nm and emitted light between 597-630 nm was recorded. For visualization of the tissue structure autofluorescence was detected at 530-595 nm by excitation of the specimen at 488 nm. Images were further analyzed using the FIJI distribution of ImageJ. To remove nonspecific detector noise, the mean fluorescence intensity of the mean fluorescence intensity of the darkest 0.1% pixels had been subtracted from all pixels in each image. Afterwards, spherical RIPK3-positive immune cells were removed from the image. The image was blurred using a median of a rolling ball radius of 8 pixels. Brightest regions were thresholded by applying the MaxEntropy auto threshold and spherical objects of 7-500 px^2^ and a circularity of 0.1-1.0 were outlined as regions of interest (ROIs). The ROIs had been manually confirmed to be immune cells and then enlarged by 0.6 µm to fully cover the RIPK3 positive immune cells. The selected ROIs were then removed from the image. The liver sinusoidal endothelium surrounds the sinusoids and is typically located at the border of the strong tissue autofluorescence. The sinusoids were detected from inverted images of the autofluorescence using the Huang auto-threshold in FIJI. Sinusoidal segments were well covered applying the FIJI particle analyzer (size: 10-2000 px^2^, circularity: 0.10-1.00). A band of 2 µm in width was then placed along the border of the sinusoidal ROIs to cover the liver sinusoidal endothelial cells (LSECs). Covering of LSECs was visually confirmed and the mean intensity RIPK3 fluorescence computed for each ROI in each image. LSEC ROIs were then removed from the image. Finally, hepatocellular RIPK3 structures appearing as small vesicles are detected in the remaining tissue (high autofluorescence) using an Otsu auto-threshold and the particle analyzer plugin (size: 0.3-15 px^2^, no limit on circularity). The results were manually confirmed. RIPK3 mean fluorescence intensity was calculated from all detected RIPK3 ROIs in each image. For both structures (LSEC and hepatocytes) one mean fluorescence intensity was calculated and is visualized in bar plots.

### Quantification of Bile Acids by Mass Spectrometry

Using a LC-MS/MS in-house assay concentration of 15 bile acids was determined in HepG2 cells and isolated primary murine hepatocytes with two different sample preparations. First, 3-fold (w/v) ethanol-phosphate-buffer (15% 0.01 mol L^-1^ phosphate buffer solution pH 7.5, 85% ethanol) was added to pre-weighed isolated primary murine hepatocytes. Samples were homogenized in a pebble mill (QiaShredder) and centrifuged at 16000 g for 5 min. The supernatant was used for bile acid quantification. Second, 1×10^6^ HepG2 cells were seeded and incubated for two days. Afterwards, cells were washed two times with 500 µL PBS, trypsinized with 100 µL 0.05% trypsin-EDTA for 3 min at 37°C and 5% CO_2_ and incubated with 1 mL DMEM to stop the trypsin reaction. The cell suspension was then centrifuged for 3 min at 4°C and 260 g. Afterwards the cell pellet was washed by resuspending it twice in 500 µL 4°C cold PBS and centrifuging for 3 min at 4°C and 260 g. The washed pellet was then resuspended in 100 µL PBS and homogenized in a pebble mill (QiaShredder). After centrifugation for 5 min at 4°C and 13000 g, the supernatant was used for bile acid quantification. The sample preparation was then followed by protein precipitation and filtration of the samples. For quantification an Agilent 1200 high performance liquid chromatography system (Agilent Technologies GmbH, Germany) with a CTC-PAL autosampler coupled to an API 4000 Triple Quadrupole mass spectrometer with electrospray ionization source (AB Sciex, Germany) was used. All chromatographic separations were performed with a reverse-phase analytical column. The mobile phase consisted of water and methanol, both containing formic acid and ammonia acetate, at a total flow rate of 300 µL min^-1^.

### Calf Intestinal Phosphatase Assay

20 µg protein lysate was incubated in calf intestinal phosphatase (CIP)-buffer (100 mmol L^-1^ NaCl, 50 mmol L^-1^ Tris-HCl, 10 mmol L^-1^ MgCl_2_, 1 mmol L^-1^ DTT, inhibitor of protease) containing 1 U CIP per µg protein for 1 h at 37°C. Afterwards, a western blot was performed with 4 µg protein as indicated in the section ‘Western Blot’.

### Lambda Phosphatase Assay

20 µg protein lysate was incubated with lambda phosphatase according to the manufacturer’s protocol (New England Biolabs). Afterwards, a western blot was performed with 15 µg protein as indicated in the section ‘Western Blot’.

### Site Directed Mutagenesis

The Q5 Site-Directed Mutagenesis PCR reaction was performed on 20 ng of the RIPK3 vector according to the supplier’s instruction (New England Biolabs) using primers and annealing temperatures (T_m_) given in **Tab. S1**. The PCR product was mixed with the provided kinase, ligase DpnI mix and transformed into NEB 5-alpha competent *E. coli*. The transformed *E. coli* were spread onto LB agar plates containing ampicillin (100 µg mL^-1^). Isolated colonies were expanded into overnight cultures and the pDNA was isolated with the ZymoPURE Plasmid MiniPrep according to manufacturer’s protocol (Zymo Research). The mutation in the sequence was confirmed by sequencing (Macrogen).

### IL-8 ELISA

Transfection was performed as described in paragraph ‘Transfection’ and stimulation as described in paragraph ‘Stimulation with Bile Acids’. HepG2 cells were washed with PBS, lysed in the RIPA buffer and the protein concentration was measured with the BCA Protein Assay Macro Kit according to the manufacturer’s protocol (Serva Gelelectrophoresis GmbH). 100 µL of the total lysate were used for the IL-8 enzyme-linked immunosorbent assay (ELISA). The assay was performed using human IL-8 ELISA Max Deluxe (BioLegend) following manufacturer’s instruction. Absorbance was measured at 450 nm and 570 nm using a plate reader (EnSpire, Perkin Elmer).

### Quantitative PCR

Transfection and stimulation was performed as described in paragraph ‘Transfection’ and ‘Stimulation with Bile Acids’. RNA isolation was performed with the Direct-zol RNA MicroPrep Kit according to manufacturer’s protocol (Zymo Research). RNA concentration and purity were assessed spectrophotometrically on NanoDrop 2000c (ThermoFisher Scientific). cDNA was generated using the RevertAid First Strand cDNA Synthesis Kit according to the manufacturer’s protocol (ThermoFisher Scientific) from 500 ng mRNA. A quantitative PCR was set up with 25 ng cDNA template, 0.5 µmol L^-1^ of each, forward and reverse primer for the individual genes of interest (Eurofins Genomics, Germany) and GoTaq qPCR 2x Master Mix (Promega, Germany). The qPCR was carried out on a RotorGene Q (Qiagen) using the following temperature protocol: 95°C for 2 min, followed by 40 cycles at 95°C for 15 s, 60°C for 60 s and 70°C for 30 s. A final ramp from 70 to 95°C for around 90 s was set to determine melting curves. The amount of mRNA for IL-8 in each sample was normalized to the sample specific Ct-value of the housekeeping gene HPRT. Fold induction of IL-8 gene expression was calculated using the Pfaffl method. [42] Primers used are found in **Table S1**.

## Results

### RIPK3 promoter methylation protects hepatocytes from necroptosis under physiological conditions

Important cellular signaling events during necroptosis are the phosphorylation of RIPK1, RIPK3 and MLKL which ultimately form an auto-lysing pore-complex. Previous studies discussed the presence of necroptosis signaling and particularly the expression of RIPK3 in liver tissue versus hepatocytes in health and liver diseases with different results. [18, 43-45] In the hepatocellular carcinoma cell line HepG2 as well as primary human hepatocytes (pHep) RIPK3 was not expressed (**Fig. 1A, Fig. S1**), which had been suggested previously from investigations on murine liver tissue and liver cell lines. [46] As expected primary human macrophages (pMФ), which served as a positive control in this experiment, expressed high levels of RIPK3 (**Fig. 1A**). [47-49]

To analyze the purity of the isolated primary hepatocytes and macrophages, a western blot against CD68 and albumin was performed from the same lysates. CD68, a protein highly expressed by monocytes and macrophages, thereby served as a macrophage specific marker whereas albumin, which has great abundance in hepatocytes, was used to characterize hepatocytes. We did not detect any expression of CD68 in the hepatocyte lysate, indicating a high purity of the samples (**Fig. 1B**). A low expression of albumin, however, was found in macrophages (**Fig. 1B**). This may be attributed to the cultivation medium that is bound to the macrophages despite washing steps or is the result of low expressed (advanced glycation end-product)-albumin in macrophages. [50]

While RIPK3 expression was absent in hepatocytes, RIPK1 and MLKL were highly expressed in three cell types HepG2, pHep, and pMФ (**Fig. 1C**). Activation of necroptosis signaling in pMФ is often achieved using TNF-α, the pan-caspase inhibitor zVAD-fmk and an inhibitor of the inhibitor of apoptosis protein (IAP) family. As previously described, this stimulation induces expression and phosphorylation of the necroptotic genes (e.g. RIPK1) (**Fig. 1C**). [51] After 6 h stimulation, pRIPK3 and pMLKL levels were elevated in pMФ, indicating initiation and propagation of necroptosis (**Fig. 1C**). [52] It is known that hepatocytes are susceptible to cytokine signaling and express TNFR1. [53] However, after 6 h of stimulation with the necroptosis inducing TNF-α/zVAD-fmk/IAP (TBZ)-mix only RIPK1 expression in the pHep increased. [54] pRIPK1, pRIPK3 and pMLKL were not detected while MLKL expression remained stable (**Fig. 1C**). Thus, in comparison to pMФ, pHep and HepG2 cell line lack an important condition to form MLKL-dependent pores and undergo necroptosis under physiological conditions after stimulation. As all signaling molecules necessary for necroptosis were present other than RIPK3, we performed DNA methylation-specific sequencing analysis of predicted CpG islands in human *RIPK3* promoter regions to elucidate a cause for this absence (**Fig. 1D**). Eight promoter elements located at the 5’ untranslated region (UTR) and initiation site of transcription of the RIPK3 gene were analyzed, located approximately −65 to +89 base pairs around the transcriptional start site. All analyzed regions in pHep were hypermethylated with relative methylated cytosine levels ranging from 37%-83% and a global methylation level over all analyzed areas of 65% (**Fig. 1D**). Similarly, HepG2 cells also showed hypermethylation in the promoter region of the RIPK3 gene. Yet, with a relative methylated cytosine amount of 21%-50% (and 35% globally), the hypermethylation is less pronounced in HepG2 cells than in pHep (**Fig. 1D**) and reflects the previous observation of global hypermethylation loss in hepatocellular carcinoma. [55] Nevertheless, the hypermethylation in HepG2 was still sufficient to suppress RIPK3 protein expression (**Fig. 1D**). In contrast, the same regions in primary pMФ expressing RIPK3 showed a relative methylation of 0%-10% and only 4% globally (**Fig. 1D**). Thus, hepatocytes and HepG2 cells have a hypermethylated RIPK3 promoter silencing protein expression, preventing these cells from undergoing necroptosis under physiological conditions (**Fig. 1D**).

### Hepatocytes express RIPK3 under pathological conditions

To investigate the RIPK3 expression in liver diseases different murine models of liver injury were utilized (**Tab. S2**) and the expression of RIPK3 in comparison to human liver diseases was analyzed (**Fig. 2**). We chose the bile duct ligation model (BDL), which resembles chronic cholestasis as a consequence of a post-hepatic, sterile injury in many ways. [56] Further, we induced warm ischemia reperfusion injury (IR) as it is a common model with pericentral cell death and injury due to generation of oxidative stress and production of inflammatory cytokines and chemokines. [57] The model is used to mimic the pathophysiological events occurring during liver transplantation. [58] The acetaminophen induced liver injury (APAP) is an acute model of liver damage that mimics a drug induced liver injury that is a common adverse effect encountered in clinical practice. [59, 60] Lastly, the peritoneal contamination and infection model (PCI) is employed to induce a murine model of sepsis that is one of the leading causes of death in intensive care units. This model results in acute peritonitis associated with a pro-inflammatory condition. [61] Liver diseases significantly vary in their etiology and pathomechanism. To gain insight into the expression regulation and relevance of RIPK3 signaling, we compared various models highlighting different aspects of chronic and acute liver failure in vivo.

**Fig. 2:**
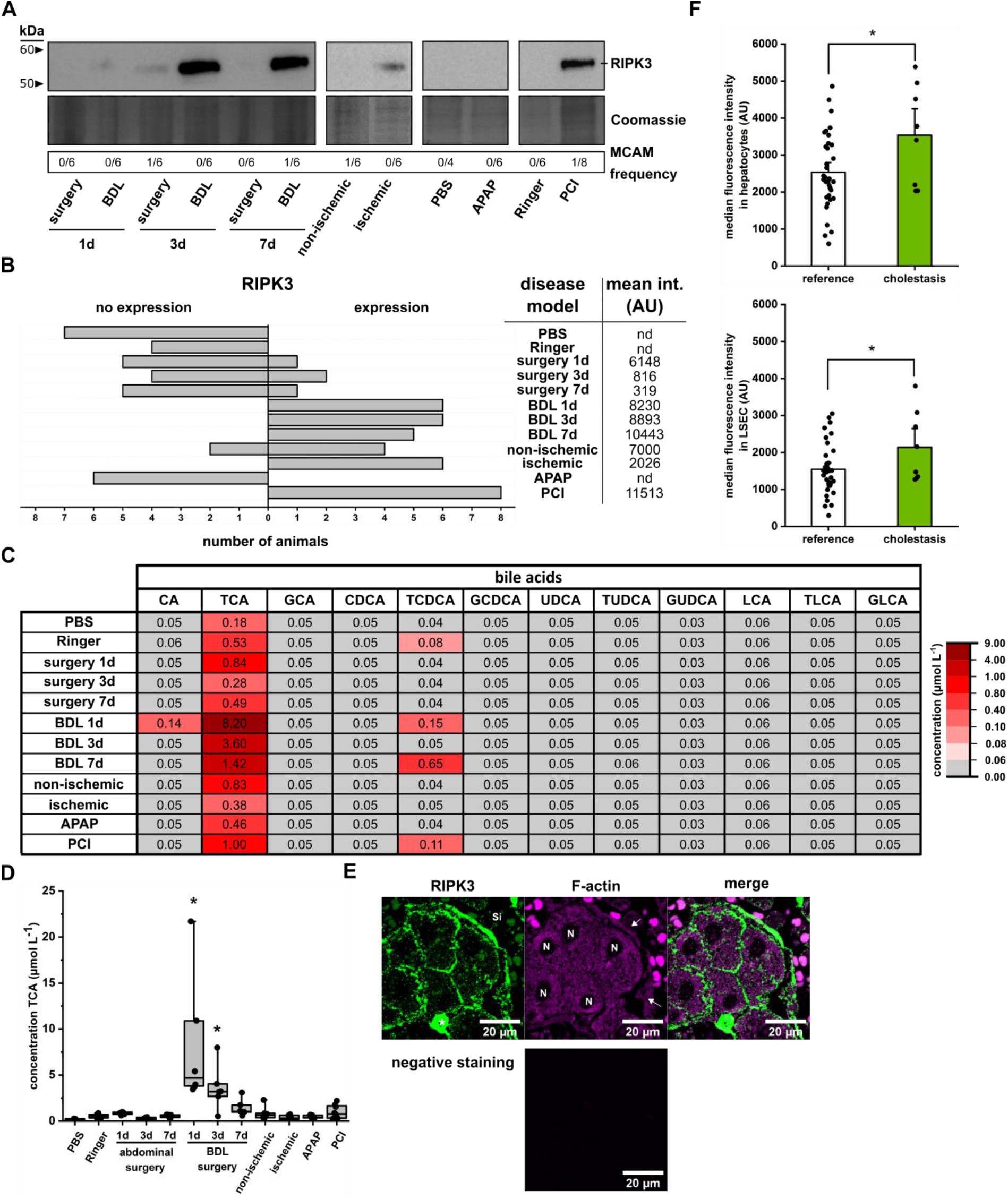
RIPK3 expression in hepatocytes is present under non-physiological conditions. (**A**) Representative western blot analysis of RIPK3 expression in different animal models of liver injury. (**B**) Absolut number of animals expressing RIPK3 and mean intensity of the signal. Control groups received Ringer acetate or PBS i.p.. Animals in the abdominal surgery group get an exposure of the liver and bile duct without additional intervention whereas the BDL group has a ligated bile duct. For the ischemic animals the left lateral branch of the portal vein was clamped to interrupt the blood flow to the left lateral lobe. Animals in the PCI and APAP group received a defined amount of human feces (PCI, 60-70 µL feces suspension) or acetaminophen (APAP, 300 mg kg^-1^ body weight) i.p.. Data are presented as a relative frequency bar plot as well as the mean intensity of the signal. (**C**) Mean concentration (µmol L^-1^) of bile acids in the isolated primary murine hepatocytes from diseased animals detected by LC-MS/MS. The LLOQ was assumed for when a bile acid was not detected. (**D**) Concentration of TCA in the different animal models used. Data are presented as a median box plot (IQR: perc 25.75, median line, whiskers from minimum to maximum). * p<0.05 vs. PBS, Kruskal-Wallis test with post-hoc Dunn’s test. (**E**) Representative fluorescence images of human liver sections stained for RIPK3 (green). The tissue was visualized by counterstaining F-actin with phalloidin (magenta). The negative staining control was treated and processed as the images, but not incubated with primary antibodies or phalloidin. Si: sinusoid, N: nucleus, *: immune cells, arrow: liver sinusoidal endothelial cells (LSEC). (**F**) Image analysis of RIPK3 expression in hepatocytes and LSECs. Data are presented as mean bar plots with standard errors and individual data-points. * p<0.05 vs. control, unpaired t-test.

We isolated hepatocytes from all models and analyzed the purity and expression of RIPK3. The purity of each individual isolated mouse hepatocyte lysate was assessed by a western blot against the melanoma cell adhesion molecule (MCAM) also known as CD146 which is highly expressed in (liver sinusoidal) endothelial cells (LSEC) [62-64] as well as various immune cells (e.g. lymphocytes, macrophages) [65-67]. In most samples MCAM was not detected (**Fig. S2**) and only a few lysates (3 out of 72) we found contamination with MCAM expressing cells. The amount of samples contaminated for each individual group is depicted as “MCAM frequency” in **Figure 2A**.

In hepatocyte lysates obtained from animals after 24 h BDL and a systemic infection (PCI), RIPK3 was highly expressed. Also, RIPK3 was detectable in the hepatocytes from the ischemic lobe of all animals after IR and in circa 60% of the respective non-ischemic lobes (**Fig. 2B**). In contrast, only a low number of animals depicted a RIPK3 expression after abdominal surgery. Further, the RIPK3 expression levels found in this model was the lowest compared to the other disease models (**Fig. 2B**). In the acutely toxic APAP liver injury model, as well as in the control groups receiving intraperitoneal injections of sterile PBS and Ringer acetate, RIPK3 expression was not detectable in isolated hepatocytes after 24 h (**Fig. 2A, Fig. 2B**). All individual results of RIPK3 expression in the primary murine hepatocytes are shown in **Figure S3**. These data demonstrate that under specific pathological conditions hepatocytes regulate RIPK3 expression and may undergo necroptosis. Further, the data suggest that this effect is more pronounced in chronic and cholestatic liver injury than in acute diseases. We then analyzed the bile acid composition in primary murine hepatocytes from the same animals which had been subjected to different models of liver injury. Consistent with literature reports, we identified taurine-conjugated CA (TCA) as the predominant bile acid in primary murine hepatocytes. [30] Taurine-conjugated primary bile acids (TCA > TCDCA > TUDCA and TLCA) were the principle accumulated bile acids in our varied models of liver injury whereas unconjugated as well as glycine-conjugated bile acids were mostly below the lower limit of quantitation (LLOQ) (**Fig. 2C**). TCA concentration increased slightly in bacterially infected (PCI) animals whereas the animals with bile duct ligation (BDL) shows a significant increase compared to the PBS control group (**Fig. 2D**). The absence of unconjugated primary bile acids shows that the liver function of biotransformation and conjugation is functional whereas hepatic clearance of bile acids is decreased. This was seen before in a study analyzing bile acid composition in liver cirrhotic patients where TCA was the most changed bile acid in the liver. [68, 69]

We then analyzed the RIPK3 expression in pathological liver sections from patients suffering chronic cholestasis. All patients included in this study underwent liver resections as part of a tumor surgery. The different underlying diseases are summarized in **Table S3**. In addition to liver sections, alanine aminotransferase (ALAT), aspartate aminotransferase (ASAT), albumin and C reactive protein (CRP) had been assessed. Patients with a total bilirubin plasma level of ≥ 21 µmol L^-1^ were considered cholestatic, while patients from this cohort with a plasma bilirubin of < 21 µmol L^-1^ served as reference. In the control group, specimens from 8 female and 10 male patients with a mean age of 63 years (0.95 confidence interval (CI): 5.9) were included. The “cholestasis group” consisted of specimens of 1 female and 7 male patients with a mean age of 61 years (CI: 11.1) (**Tab. S3**). Along with bilirubin, the other markers of liver damage and inflammation assessed (ALAT, ASAT, albumin and CRP) were significantly elevated in the cholestasis group (**Tab S3**).

It is of note that RIPK3 expression was found in hepatocytes, LSECs and immune cells of all specimens regardless of their bilirubin level, indicating that the RIPK3 expression is induced in hepatocytes upon various liver injuries (**Fig. 2E**). In all groups the mean RIPK3 fluorescence intensity (FI) in LSECs was lower than in hepatocytes. Further, the mean RIPK3 FI was significantly increased in specimens from patients suffering hyperbilirubinemia compared to the reference group. This effect was present in both cell types, but more pronounced in hepatocytes compared to LSECs (**Fig. 2F**).

### Bile acids are sensitive to affect RIPK3 activation that induces inflammation

After confirming the expression of RIPK3 in cholestatic liver diseases we sought further the effects of bile acids on the activation of RIPK3 and subsequent signaling events. Therefore we overexpressed a previously characterized human RIPK3 (NM_006871.3) construct with an N-terminal FLAG-tag driven by a CMV promoter (RIPK3-FLAG, [39]) in HepG2 cells, that do not express RIPK3 endogenously (**Fig. 3A**). The transfection with the vector backbone alone did not lead to an expression of RIPK3, thus the detected RIPK3 is fully attributed to the RIPK3-FLAG construct in the expression vector (**Fig. 3A**). Further it is of note that the basal amount of bile acids in these HepG2 cells were mostly not detectable (nd) and therefore results with external bile acid stimulation may be attributed to the different bile acids supplemented for stimulation (**Fig. 3B**). We incubated HepG2 cells with bile acids at 50 µmol L^-1^ dissolved in methanol, a non-physiological concentration reported to be present in human liver tissue during cholestasis, [70], which they may take up by specialized transporters such as organic anion transporter (pumps) or through passive diffusion. [71] LCA and glycine-LCA (GLCA) exhibited cell toxicity and caused significant membrane damage at 50 µmol L^-1^, as examined by quantifying lactate dehydrogenase in the supernatant (**Supplementary Methods**), and were therefore used at their highest non-toxic concentration for stimulation (LCA: 5 µmol L^-1^, GLCA: 10 µmol L^-1^) (**Fig. S4**).

**Fig. 3:**
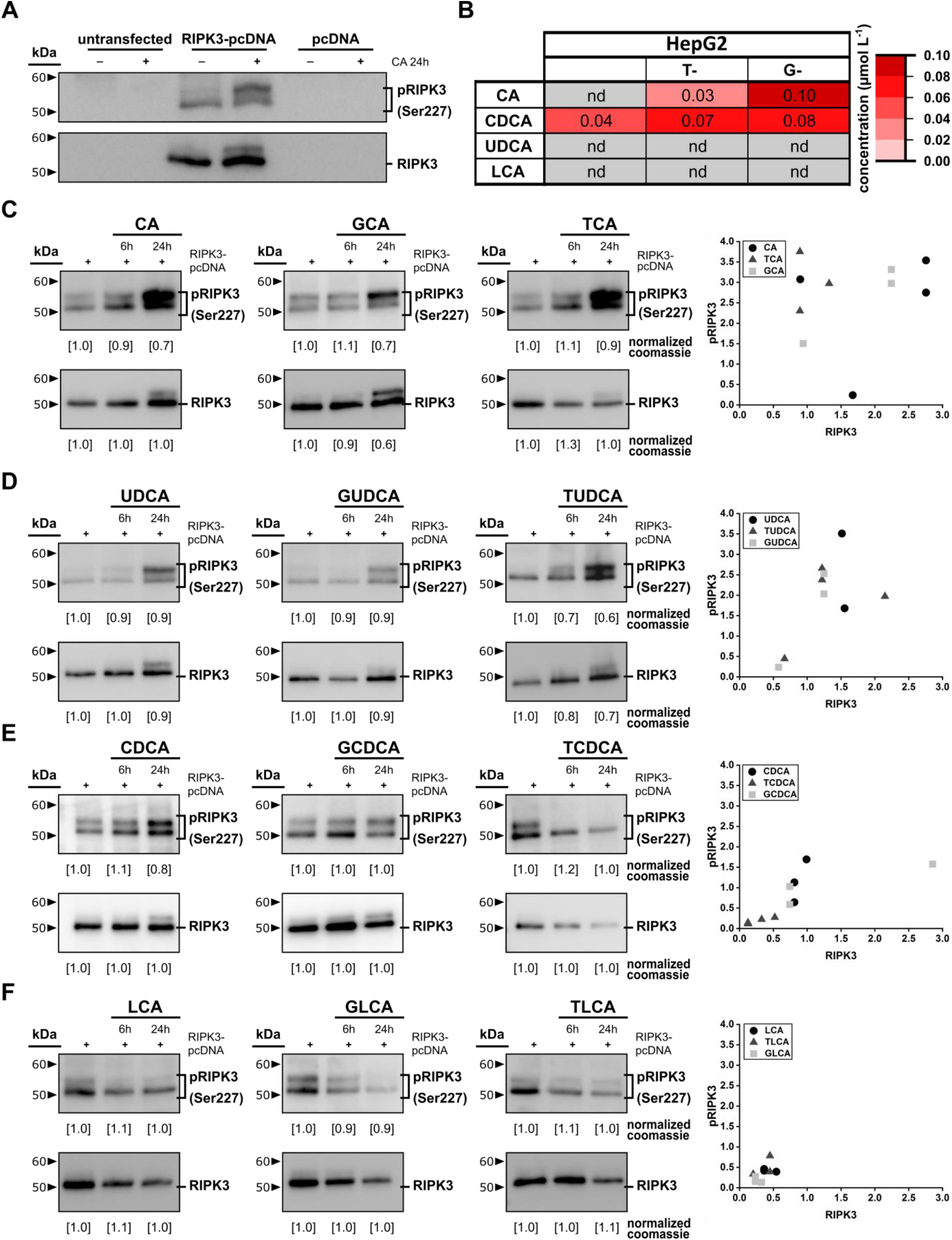
Bile acids are able to affect RIPK3 activation. (**A**) A representative blot of RIPK3 expression and phosphorylation after transfection of pcDNA plasmid and RIPK3-pcDNA plasmid. (**B**) Amount of bile acids in the hepatocellular carcinoma cell line (HepG2) detected by LC-MS/MS. The analysis was performed on 3 individual samples. Grey colored values were not detected (nd). (**C-F**) RIPK3 phosphorylation and expression upon stimulation with different endogenous bile acids for 6 or 24 h. Stimulation with un-, taurine (T)- or glycine (G)-conjugated (**C**) cholic acid (CA), (**D**) ursodeoxycholic acid (UDCA), (**E**) chenodeoxycholic acid (CDCA) and (**F**) lithocholic acid (LCA). (**C-F**) Coomassie staining of SDS-PAGE gel was used as a loading control. n=3-4.

Overexpression of RIPK3-FLAG leads to a basal phosphorylation in HepG2 cells (**Fig. 3A, Fig. 3C-F**), suggesting that various endogenous mechanisms may trigger its activation. The unconjugated hydrophilic bile acids CA and UDCA increased RIPK3-FLAG phosphorylation and RIPK3-FLAG expression itself after 6 and 24 h (**Fig. 3C, Fig. 3D**). Also, CDCA shows a minor effect on the RIPK3-FLAG phosphorylation while not affecting the RIPK3-FLAG expression (**Fig. 3E**). In contrast LCA decreased both, RIPK3-FLAG phosphorylation and expression within 24 h (**Fig. 3F**). TCA, glycine-conjugated CA (GCA) as well as glycine- and taurine-conjugated UDCA (TUDCA and GUDCA) reduced the stimulatory effects of RIPK3-FLAG phosphorylation of CA or UDCA but did not abolish them (**Fig. 3C, Fig. 3D**). Interestingly, taurine-conjugated CDCA (TCDCA) decreased the expression and phosphorylation of RIPK3-FLAG without signs of toxicity, whereas the glycine-conjugation of CDCA (GCDCA) induces no changes (**Fig. 3D**). The effects of GLCA taurine-conjugated LCA (TLCA) are equivalent to the unconjugated form (**Fig. 3F**). These findings indicate a high complexity of the underlying metabolic signaling network affecting RIPK3 expression and necroptosis on multiple levels. In the context of the heterogeneous bile acid toxicity these effects are also well known as the bile acid paradox. [72, 73] In connection with the ability to induce RIPK3-dependent necroptosis we hypothesized that epigenetic RIPK3 suppression allows hepatocytes to resist noxious stimulation from endogenous metabolites as bile acids and avoid resulting chronic pro-inflammatory and death signaling. Hepatic synthesis and metabolism of bile acids, and perhaps other metabolites, could lead to permanent activation of RIPK3 and subsequent inflammation or even cell death. [74] As noted above, bile acids modulate the expression and phosphorylation of RIPK3. We observed, consistent with previous literature, a phospho-serine (Ser) 227 RIPK3-FLAG positive double band also detectable using the RIPK3 antibody. [8, 75] As shown by Chen 2013, RIPK3 contains multiple phosphorylation sites (Ser199, Ser227) that exhibit different functions and are both indispensable for the necroptosis induction. [75] Ser199 is the important residue for induction of kinase activity [76] whereas Ser227 is crucial for the induction of necroptosis [75]. Additionally, ubiquitination was reported to be regulated during RIPK3 activation. [77] Both post translational modifications may lead to the observed mass shift in HepG2 cells. [78] We were able to exclude the ubiquitination of RIPK3-FLAG processing in our model as a consequence for the mass shift (**Fig. S5**). To investigate a hyperphosphorylation we incubated the protein lysate with calf intestinal phosphatase (CIP) or Lambda phosphatase (LP), which possess a high specificity for phospho-serine, -threonine and -tyrosine residues. CIP and LP, both abolished the pSer227 RIPK3-FLAG signal and diminished the mass shift observed in the total RIPK3-FLAG blot (**Fig. 4A**). Thus, bile acids are not only able to modulate the RIPK3 phosphorylation but further modulate its hyperphosphorylation, which is a necessity for the activation of necroptosis signaling. [79]

**Fig. 4:**
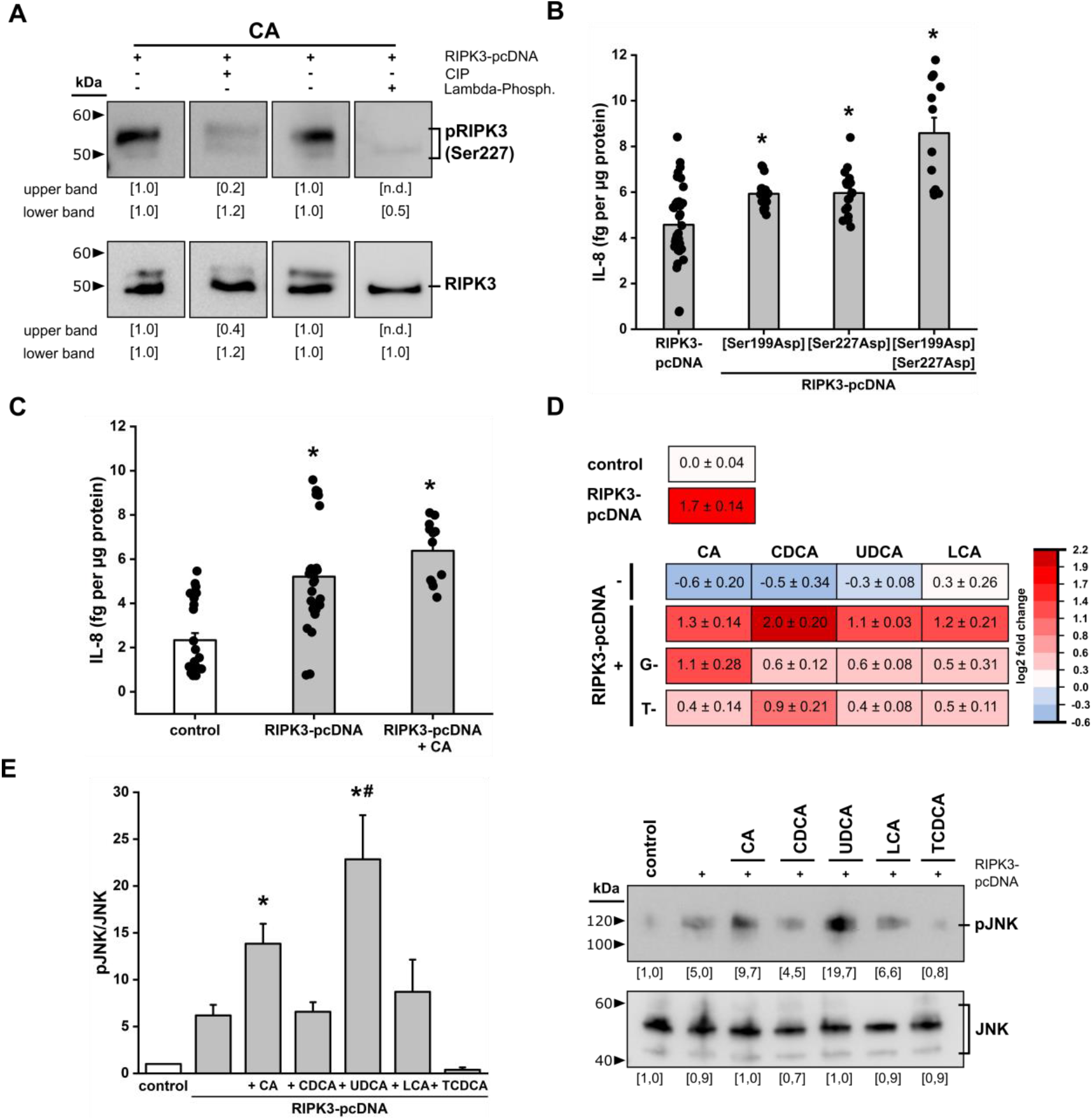
RIPK3 facilitates bile acid mediated inflammation. (**A**) Transfected and stimulated HepG2 cells are lysed and treated with calf intestinal phosphatase (CIP) for 30 min at 37°C or Lambda phosphatase for 30 min at 30°C before western blot analysis of RIPK3 expression and phosphorylation. The upper lane was not detected (nd) in the treated samples. (**B**) Secretion of the pro-inflammatory cytokine IL-8 in HepG2 cells after transfection with RIPK3-pcDNA plasmid and phospho-positive mutants of RIPK3 to aspartic acid (Asp). Data are presented as mean bar plot with standard error and individual data-points. * p<0.05 vs. RIPK3-pcDNA, Kruskal-Wallis test with post-hoc Dunn’s test. (**C**) Relative secretion of the pro-inflammatory cytokine IL-8 in HepG2 cells after transfection with RIPK3-pcDNA. Data are presented as mean bar plot with standard error and individual data-points. * p<0.05 vs. control, Kruskal-Wallis test with post-hoc Dunn’s test. (**D**) mRNA levels of IL-8 in HepG2 cells with and without transfection of the RIPK3-pcDNA and stimulation with different bile acids. Data are presented as mean with standard error in the heatmap. (**E**) JNK expression and phosphorylation upon stimulation with different endogenous bile acids for 24 h. Coomassie staining of SDS-PAGE gels was used as a loading control. Data are presented as mean bar plot with standard error. n=3. * p<0.05 vs. control and ^#^ p<0.05 vs. RIPK3-pcDNA, one-way ANOVA with post-hoc Dunnett’s test.

The bile acid dependent activation and modification of RIPK3 primes the necroptotic pathway and stimulates a phosphorylation of MLKL which then translocate to the plasma membrane and induces cell rupture followed by an inflammatory reaction. In hepatocytes, the secretion of interleukin-8 (IL-8) is specific as an early inflammatory insult to recruit immune cells, especially neutrophils, and trigger secondary tissue damage as well as repair in various diseases. [80] Therefore human hepatoma HepG2 cell line was used as a representative model of the human liver that display a high degree of morphological and functional differentiation to generate reproducible results. [81]

To mimicking a permanent activation of both important phosphorylation sites of RIPK3 (Ser199, Ser227), mutation of the serine residue to aspartic acid (Asp) promoted the IL-8 secretion significantly especially when both phosphorylation sites were mutated to activating Asp residue (**Fig. 4B**). This demonstrates that expression, activation and hyperphosphorylation of RIPK3, as under in vitro cholestatic conditions, results in a necroptosis-related inflammatory response in HepG2 cells. The IL-8 level increases significantly upon expression of RIPK3-FLAG in HepG2 cells and is further enhanced by stimulation with the bile acid CA, the bile acid with the strongest RIPK3-FLAG activating and phosphorylating function on protein level (**Fig. 4C**). This shows that the bile acid CA is able to induce the necroptotic pathway by activating RIPK3 resulting in inflammation. Besides secretion, RIPK3-FLAG expression and stimulation with different bile acids affected the gene expression of IL-8 (**Fig. 4D**). HepG2 cells that don’t express detectable levels of endogenous RIPK3 decrease the expression of IL-8 after stimulation with different unconjugated bile acids which could be interpreted as a protective effect of the bile acids. RIPK3-FLAG transfection immediately leads to a strong expression of IL-8 that correlates with the direct phosphorylation of RIPK3-FLAG seen on protein level (**Fig. 3, Fig. 4D**). Stimulation of RIPK3-FLAG expressing HepG2 cells with unconjugated bile acids hardly affected the IL-8 gene expression. Solely the hydrophobic bile acid CDCA induced IL-8 gene expression significantly (**Fig. 4D**). Strikingly, the hydrophilic bile acid CA which induced activation and phosphorylation of RIPK3-FLAG as well as secretion of IL-8 did not affect the gene expression of IL-8 (**Fig. 3C, Fig. 4C, Fig. 4D**). Incubation with all glycine- and especially taurine-conjugated bile acids however reduced the IL-8 gene expression in comparison to their unconjugated counterparts. This is attributable to the reduction of the hydrophobicity and in turn the increased cell protective properties due to increased hydrophilicity and impermeability to the cell membrane. [36, 82] As stated in previous literature it is known that IL-8 secretion may be controlled by different pathways whereby the Jun-(N)-terminal kinase (JNK) signaling pathway is one prominent example. [83, 84] Therefore we used the four primary, unconjugated bile acids (CA, CDCA, UDCA, LCA) as well as TCDCA, the taurine-conjugated form of the strongest IL-8 gene expression activator, to investigate JNK activation by phosphorylation at Thr183/ Tyr185. RIPK3-FLAG overexpression directly stimulated JNK phosphorylation that agreed with the immediately seen RIPK3-FLAG phosphorylation and increased IL-8 secretion and gene expression (**Fig. 3, Fig. 4C-E**). Stimulation with the hydrophilic bile acids (CA, UDCA), that lead to activation and phosphorylation of RIPK3-FLAG, further enhanced the phosphorylation of JNK (**Fig. 4E**). Similar to the reducing effects on RIPK3-FLAG expression and phosphorylation, TCDCA strongly suppressed JNK phosphorylation even in the presence of RIPK3-FLAG (**Fig. 4E**). These results demonstrate that RIPK3 expression and phosphorylation induces necroptosis, which in turn induces IL-8 secretion regulated by JNK. This mechanism may then lead to both: an activation of repair mechanisms or progression of the liver disease.

## Discussion

Our findings indicate that RIPK3 silencing is a hallmark mechanism within hepatocytes to avoid necroptosis, initiation of pro-inflammatory signaling, and pore formation. [85] We have identified bile acids as a class of molecules able to trigger the activation and phosphorylation of RIPK3 that in turn results in an RIPK3-dependent IL-8 response in hepatocytes, which is known to trigger local tissue remodeling and infiltration. [86] Increased IL-8 secretion and gene expression is ascribed to the activation of JNK, a pathway involved in several physiological and pathological processes. [87] Thus, the RIPK3-mediated activation of the JNK pathway in hepatocytes may not only lead to an increased local IL-8 signaling but also trigger subsequent pro-inflammatory adaptations such as cell death, cell survival and proliferation in a cell type-specific manner. [88] The JNK pathway is one of the three major groups of mitogen-activated protein kinases (MAPK) which plays a significant role in acute as well as chronic liver injuries by regulating the metabolism and cell death pathway in the liver. [89] Previous studies show that bile acids are able to activate the JNK pathway which results in an inhibition of the bile acid synthesis. [90, 91] Besides this, activation of JNK in general is also known to contribute to the expression of pro-inflammatory cytokines (IL-8, IL-6, IL-17). [84, 92, 93] Taken together, the results of this manuscript describe a hepatocytic pRIPK3-pMLKL-pJNK-IL-8 axis whereby the effect of other MAPK (p38, extracellular-signal-regulated kinase) need to be evaluated further.

In conclusion, the observed RIPK3 promoter methylation which suppresses RIPK3 expression in hepatocytes prevents metabolite (e.g. bile acid) induced hepatocellular injury under physiological conditions and may be viewed as a key-protective mechanism allowing hepatocytes to synthesize and transform metabolites that otherwise would constantly trigger inflammatory signaling in hepatocytes under physiological conditions.

Under pathophysiological conditions RIPK3 expression however was rapidly induced in mice and in all liver-related pathologies the expression of RIPK3 was found in hepatocytes and LSECs. Hepatocytes seemingly employ epigenetic RIPK3 silencing as an endogenous master switch protecting them from endobiotic metabolites, e.g. bile acids. An increase of conjugated bile acids, due to the great capacity for biotransformation and storage function of hepatocytes, is often observed first in the pathophysiology of diseases. [94-96] High concentrations of especially CA and TCA, causes RIPK3 hyperphosphorylation which is crucial for the consecutive activation of necroptosis, and inflammatory JNK-signaling. [97, 98] TCA and TUDCA are able to facilitate cell survival and act as anti-cholestatic metabolites. [99] Our results further demonstrate that the same choleretic bile acids (TCA, TUDCA) are able to induce necroptosis in hepatocytes in various liver diseases associated with a hepatocellular RIPK3 expression. This demonstrates a novel pathophysiological mechanism in the progression of liver diseases.

The hepatocellular accumulation of TCA, and in some cases also CA, during various liver injuries supports the notion that hepatocytes employ necroptosis signaling to induce tissue remodeling and inflammation. Hyperphosphorylation of RIPK3 had been recognized previously as a key event in the activation cascade of RIPK3 and necroptosis. [79] As mentioned before, Ser199 (kinase activity) and Ser227 (interaction with MLKL and induction of necroptosis) contains specific functions during the activation of RIPK3. Activation of RIPK1 induces the activation of RIPK3. The following hyperphosphorylation of RIPK3 induces the interaction with MLKL via the RIP homotypic interaction motif. The full mechanism of RIPK3 hyperphosphorylation has yet to be elucidated, but it seems a likely possibility that activation of RIPK3 induces a phosphorylation at Ser199 which in turn auto-phosphorylates Ser227 for the formation of a stable complex with MLKL. [75, 100] Hydrophobic TCDCA on the other hand, also frequently increased during cholestasis, reduces the expression and phosphorylation of RIPK3. This suggests that other unconjugated bile acids besides CA exert their cell toxicity primarily by the regulation of apoptosis and due to their function as detergent in high concentrations may lead to direct tissue necrosis. [36, 101-103] The accumulation of bile acids may not represent the primary cause of liver injury but likely promote disease progression and chronification due to a chronic inflammatory response. [70] Further, this study’ findings support previous controversial results obtained from RIPK3 knockout mice after various types of experimental liver injury. Knockout mice were especially protected from injuries that commonly result in chronic liver diseases (e.g. obstructive cholestasis, ethanol induced liver injury), inflammatory liver diseases (e.g. concanavalin-A hepatitis, fecal induced sepsis) and ischemia reperfusion damage [18, 20, 24, 104], but not in the situation of acute toxic damage as caused by e.g. acetaminophen, that may lead to an accumulation of unconjugated bile acids in blood [25, 26].

As RIPK3 knockout mice are protected from liver damage during different types of chronic injury, we postulate that the epigenetic profile, which is regulated in a highly dynamic manner during injury and liver regeneration, may be remodeled rapidly during cell stress and will modify hepatocyte susceptibility to endogenous metabolites, inflammatory signaling and cell death.

In summary, the RIPK3 expression is highly dynamic and is physiologically silenced in hepatocytes allowing them to carry out biotransformation of endogenous and exogenous substances even in high concentrations that may otherwise be toxic to the cells without triggering chronic inflammation. On the other hand, our results indicate that RIPK3 expression is rapidly increased during cholestatic liver injuries and the accumulation of bile acids, particularly hydrophilic ones (unconjugated and conjugated CA and UDCA), triggers the activation of inflammation through the necroptosis pathway. Especially in chronic situations where tissue regeneration and resolution of the injury is not achieved, the chronic activation of RIPK3 is another player contributing to the persistence of consistent liver injury.

## Limitation of the study

The cell death mechanism referred to as necroptosis attracted a great attention over the time especially due to the controversial discussion about its function in the liver. It remains to be elucidated whether RIPK3 is expressed in all types of liver diseases or just in case of obstructive cholestasis. In human specimens the investigation of RIPK3 signaling is particularly challenging due to the lack of healthy reference material. We have used bilirubin as a biomarker to depict differences, however we are aware that bilirubin rises late in the cause of liver disease and a prospective evaluation with a differential analysis of hepatocellular vs. plasma bile acids concentration may shine further light on the regulation of necroptosis signaling in these cells.

## List of Abbreviation

(RIPK3): receptor-interacting serine/threonine-protein kinase 3
(MLKL): mixed lineage kinase domain-like protein
(RIPK1): receptor-interacting serine/threonine-protein kinase 1
(TNFR1): tumor necrosis factor receptor type-1
(CASP8): caspase 8
(JNK): Jun-(N)-terminal kinase
(MAPK): mitogen-activated protein kinases
(CA): cholic acid
(GCA): glycine-conjugated CA
(TCA): taurine-conjugated CA
(CDCA): chenodeoxycholic acid
(GCDCA): glycine-conjugated CDCA
(TCDCA): taurine-conjugated CDCA
(LCA): lithocholic acid
(GLCA): glycine-conjugated LCA
(TLCA): taurine-conjugated LCA
(UDCA): ursodeoxycholic acid
(GUDCA): glycine-conjugated UDCA
(TUDCA): taurine-conjugated UDCA
(pHep): primary human hepatocytes
(pMФ): primary human macrophages
(LSEC): liver sinusoidal endothelial cells
(FCS): fetal calf serum
(LDM): liver digest medium
(CIP): calf intestinal phosphatase
(LP): Lambda phosphatase
(Ser): serine
(Asp): aspartic acid
(Ala): alanine
(Arg): arginine
(BDL): bile duct ligation
(IR): ischemia reperfusion injury
(APAP): acetaminophen induced liver injury
(PCI): peritoneal contamination and infection
(UTR): untranslated region
(MCAM): melanoma cell adhesion molecule
(ALAT): alanine aminotransferase
(ASAT): aspartate aminotransferase
(CRP): C-reactive protein
(CI): confidence interval
(FI): fluorescence intensity
(nd): not detectable
(LLOQ): lower limit of quantitation

## Author contribution

**JH** designed, carried out and analyzed experiments, **LX, JH** and **ATP** planned and conducted animal experiments and cell isolation, **TK** isolated primary macrophages and analyzed the protein expression in primary hepatocytes and primary macrophages, **SN** measured bile acids in primary human hepatocytes and HepG2 cells, **JM** analyzed the gene expression in HepG2 cells, **MG** isolated and differentiated peripheral blood mononuclear cells, **SD** and **ATP** supervised experiments and analysis, **NG** planned and supervised the experiments on human specimens, **MB** and **ATP** guided the study, **JH** and **ATP** drafted the manuscript and prepared figures. **All authors** contributed to writing sections of the manuscript.

## Acknowledgement

We acknowledge Prof. Mervyn Singer, Bloomsbury Institute of Intensive Care Medicine, United Kingdom, for the critical discussions and correction of the manuscript. We thank Dustin Beyer (Friedrich Schiller University Jena) and Felix Biedermann (Friedrich Schiller University Jena) for their support with animal experimentation and data collection. Tissue sections were kindly provided by Prof. Dr. Amelie Lupp (Institute of Pharmacology and Toxicology). Dr. Oliver Sommerfeld (Department of Anesthesiology and Intensive Care Medicine) provided clinical and demographic metadata.

## Introductory Statement

Hepatocytes suppress RIPK3 expression to protect themselves from bile acid induced inflammation and necroptosis. Expression of RIPK3 is induced in chronic liver diseases rendering hepatocytes particularly vulnerable to cell death.

## Tables

**Table 1:**
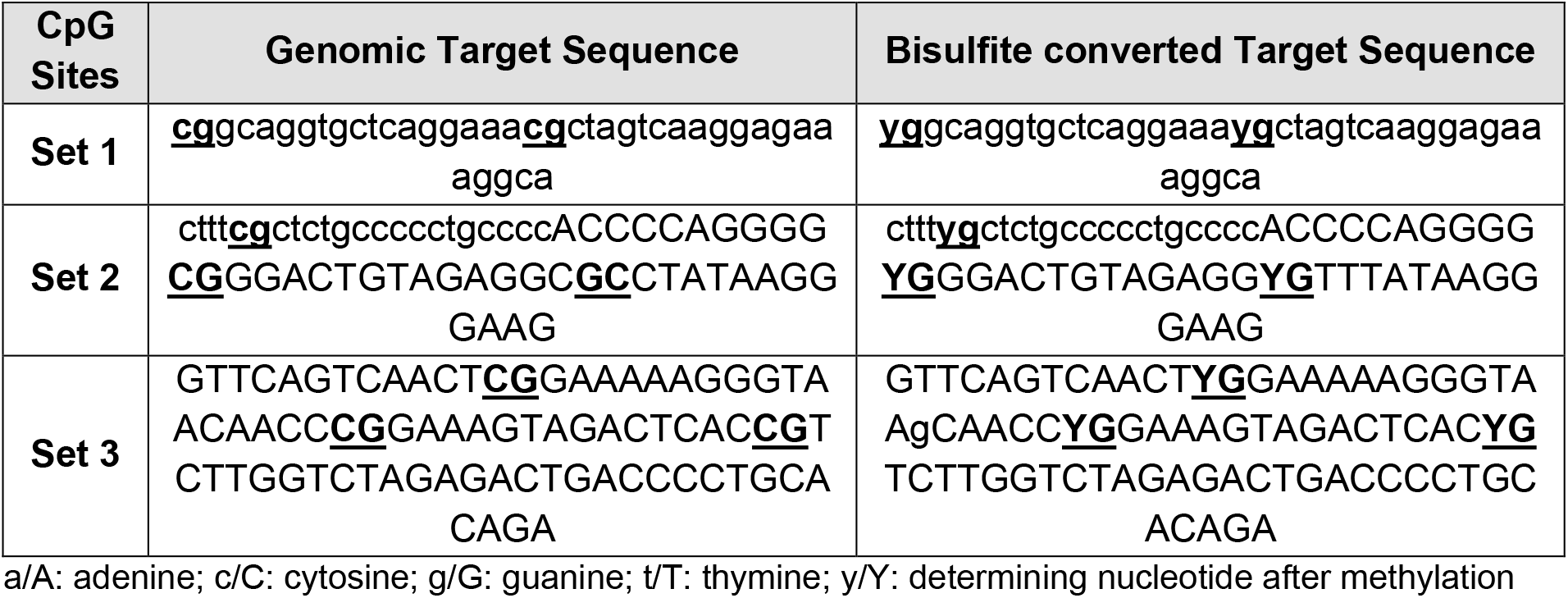
Sequences of 8 representative CpG sites in the methylation analysis. Lowercase letters represent the sequence upstream of the 5’ UTR region and capital letters the sequence in the 5’ UTR region.

## Supplemental Methods

### LDH Assay

Lactate dehydrogenase (LDH) activity can be used as a marker of cell membrane integrity to assess the cytotoxicity caused by compounds. For measuring the effects of the bile acids LCA, GLCA and TLCA on HepG2 cells the CytoTox 96 Non-Radioactive Cytotoxicity Assay (#G1780, Promega), was performed. Following incubation with the bile acids as described in ‘Stimulation with Bile Acids’, 50 µL of cell supernatant were transferred into a 96-well plate and mixed with 50 µL of the substrate. After 30 min incubation the stop solution was added and absorbance was measured at 490 nm. The LDH release was determined by subtracting the media background and calculating the amount according to positive control.

### Cell Culture

Cryopreserved primary human hepatocytes donor pools (pHep) with 20 male (average age: 36.9 years, average BMI: 26.3) or 20 female (average age: 41.1 years, average BMI: 29.2) donors were purchased from Lonza, Switzerland. Experiments with the cells were performed as indicated in the section ‘Western Blot’.

## Supplementary Figures

**Supplementary Fig. S1:**
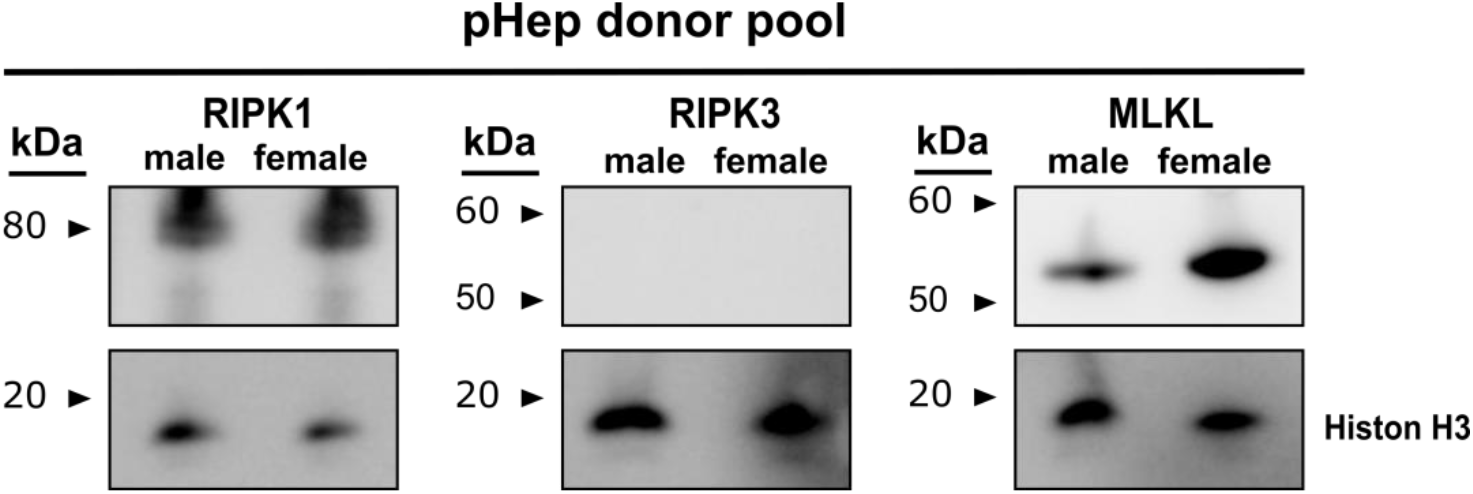
Analysis of RIPK1, RIPK3 and MLKL expression of a pHep donor pool. Protein expression of molecules in the necroptosis signaling pathway detected in male and female pHep donor pool (20 donors each). 20 µg proteins were loaded. Histone H3 was used as a loading control.

**Supplementary Fig. S2:**
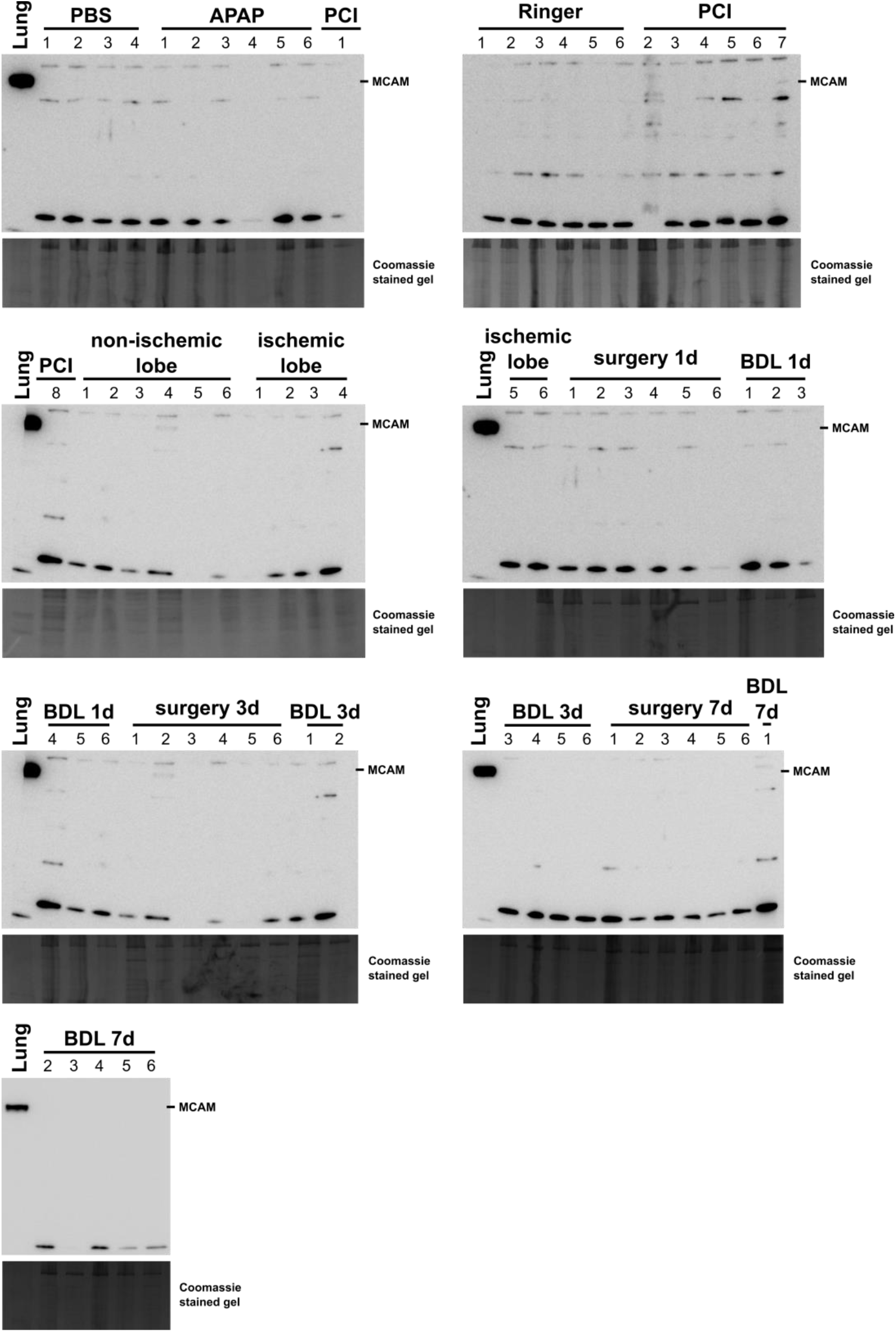
Purity of primary murine hepatocytes. Protein expression of the endothelia cell marker melanoma cell adhesion molecule (MCAM/ CD146) in different primary murine hepatocytes was analyzed by western blot. Lung lysate was used as positive control. 10 µg proteins were loaded. Coomassie stained gels were used as loading control.

**Supplementary Fig. S3:**
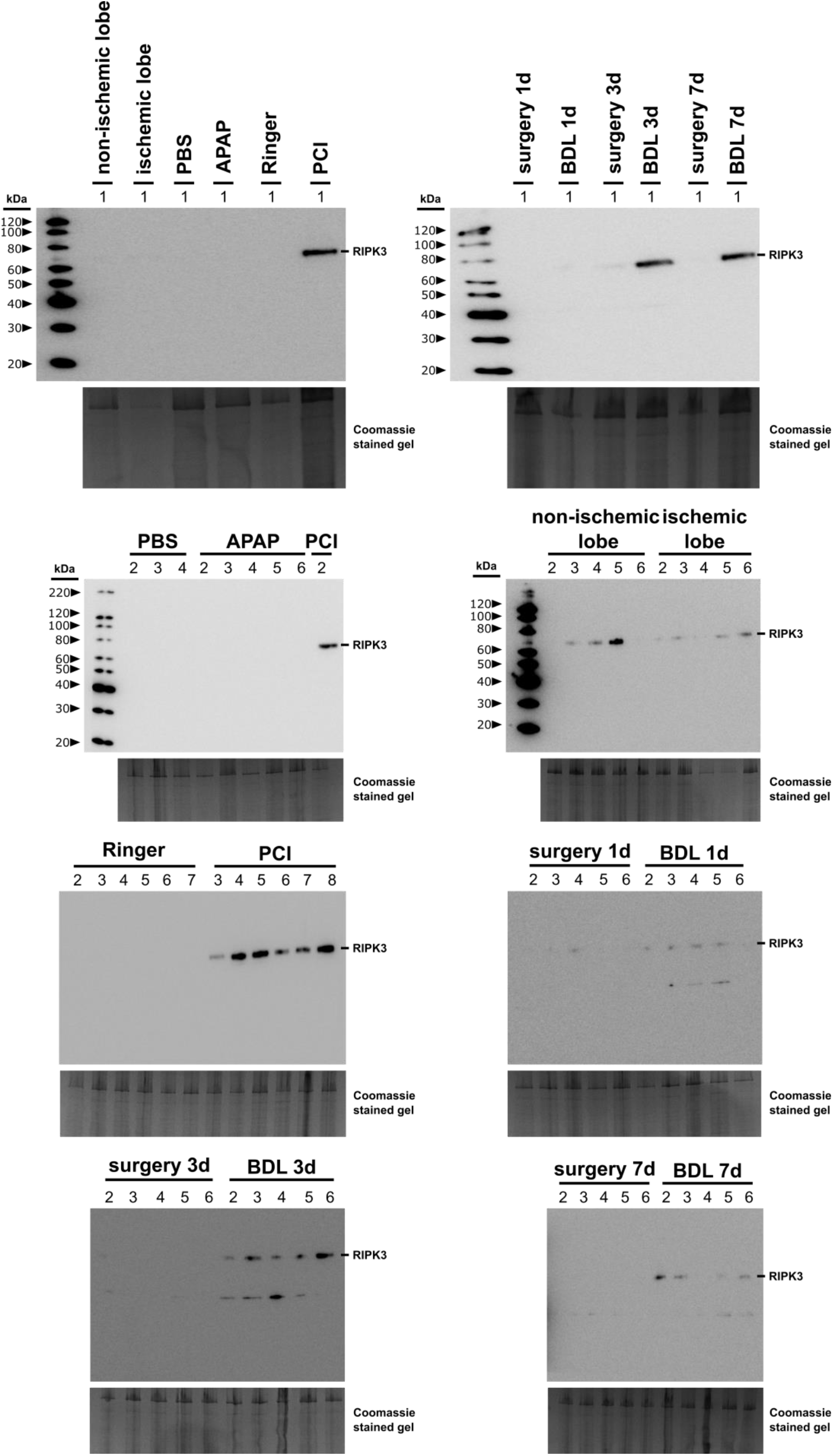
RIPK3 Expression in primary murine hepatocytes. Protein expression of RIPK3 in different primary murine hepatocytes was analyzed by western blot. 10 µg proteins were loaded. Coomassie stained gels were used as loading control.

**Supplementary Fig. S4:**
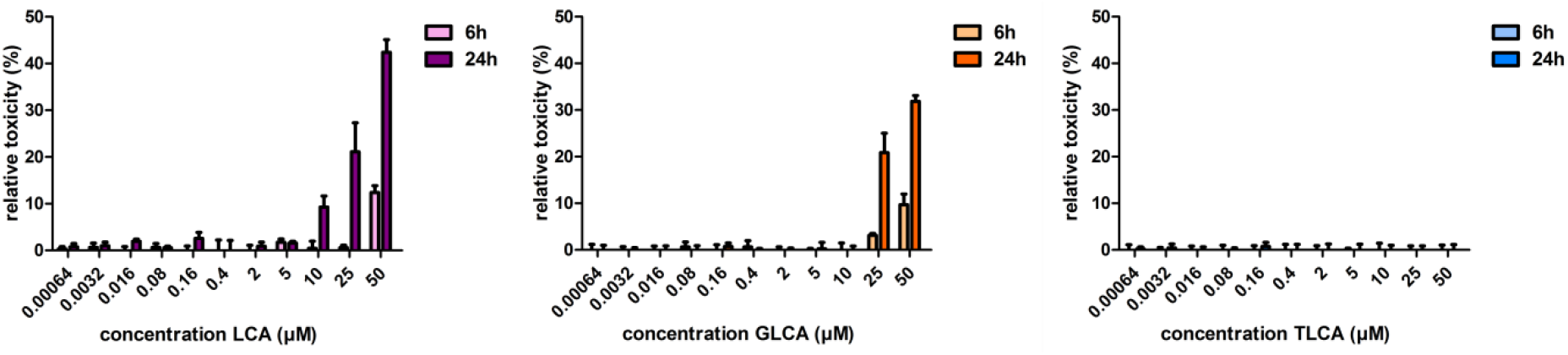
Cytotoxicity of (un)conjugated lithocholic acid. HepG2 cells were exposed to lithocholic acid (LCA) and its glycine- (GLCA) and taurine- (TLCA) conjugated metabolites for 24 h in defined concentrations. Toxicity was investigated by the LDH-assay. Data are presented as mean bar plot with standard error after subtracting the control. n=4.

**Supplementary Fig. S5:**
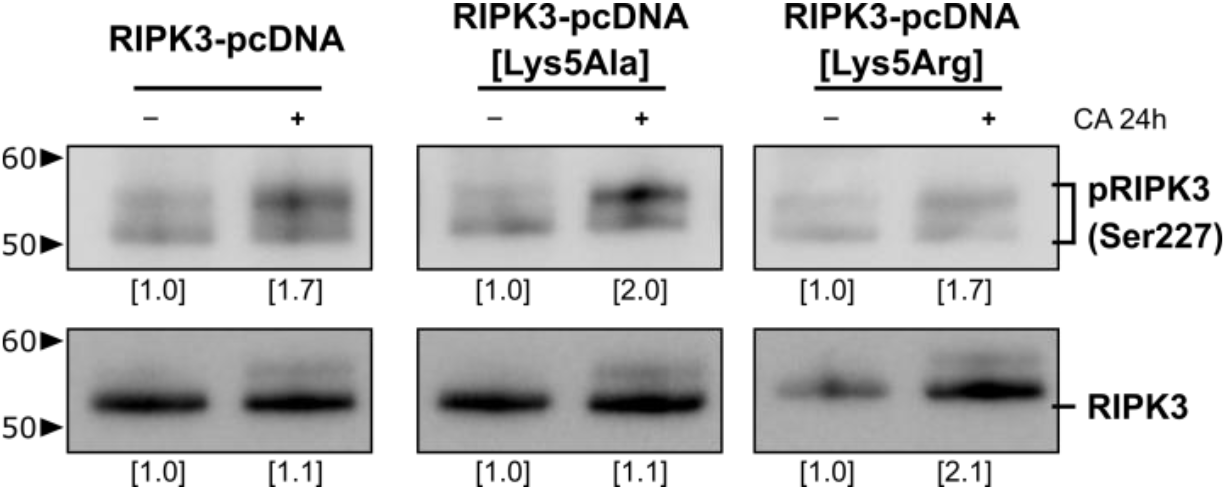
Analysis of ubiquitination induced RIPK3 band mass shift. RIPK3 expression and phosphorylation in HepG2 cells transfected with a mutated plasmid at the ubiquitination site Lys5 to alanine (Ala, permanent inhibition) and arginine (Arg, permanent activation) was analyzed by western blot. n=1.

## Supplementary Tables

**Supplementary Table S1:**
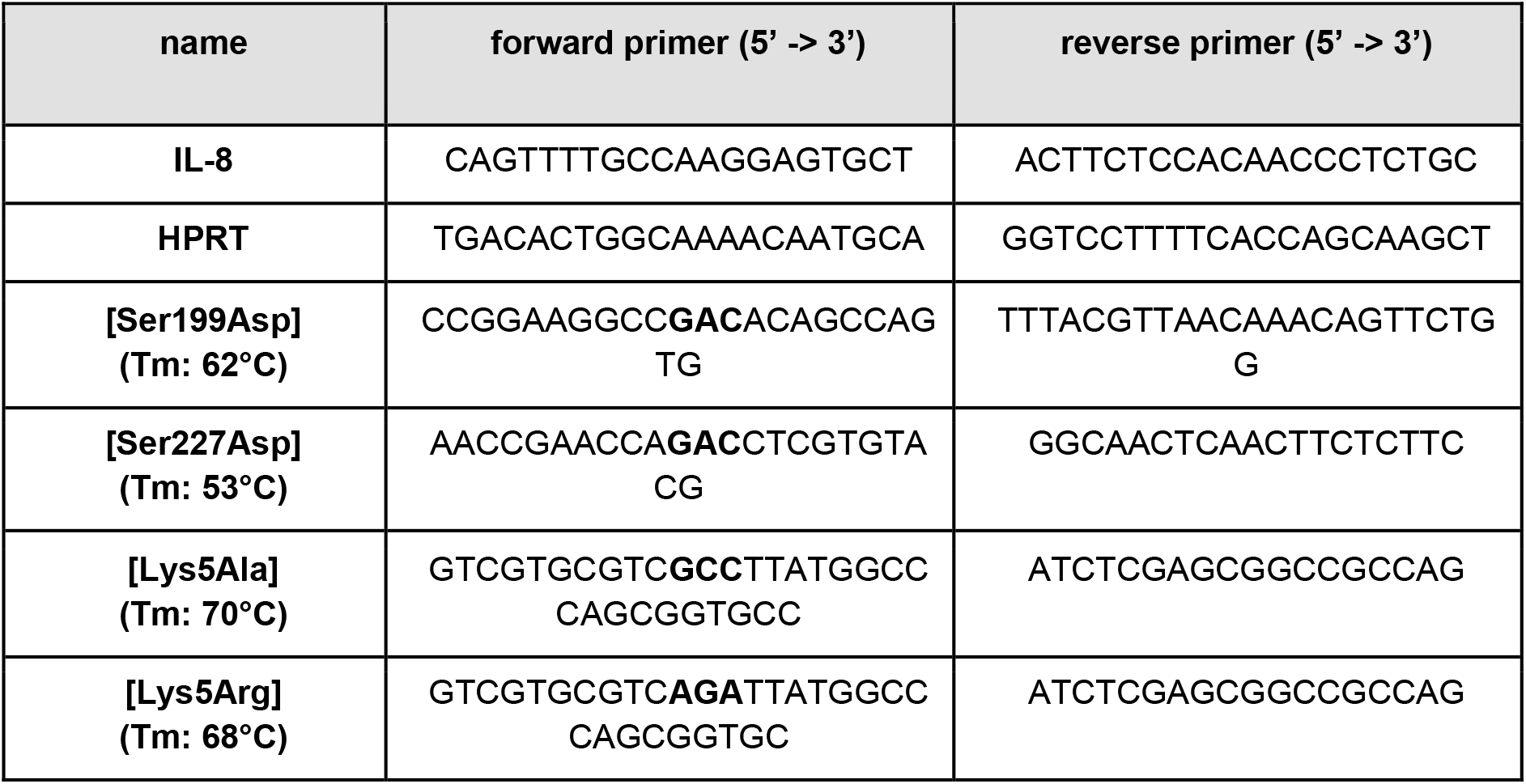
Sequences and melting temperature of used primers in the manuscript.

**Supplementary Table S2:** Overview of different liver disease parameters. mean±SD.

| | change in body weight (g) | ALAT<br>($\mu\text{mol (L*s)}^{-1}$) | ASAT<br>($\mu\text{mol (L*s)}^{-1}$) |
| --- | --- | --- | --- |
| **PBS i.p.** | 2.4 ± 0.4 | 0.9 ± 0.1 | 4.8 ± 1.1 |
| **Ringer acetate i.p.** | 0.5 ± 0.5 | 0.8 ± 0.1 | 4.4 ± 0.5 |
| **abdominal surgery 1 d** | -0.4 ± 0.7 | 1.0 ± 0.2 | 5.6 ± 0.7 |
| **abdominal surgery 3 d** | -1.0 ± 1.1 | no values | 4.6 ± 0.8 |
| **abdominal surgery 7 d** | -0.8 ± 0.7 | 0.6 ± 0.1 | 3.0 ± 0.3 |
| **BDL surgery 1 d** | -1.4 ± 0.6 | no values | no values |
| **BDL surgery 3 d** | -2.1 ± 1.4 | 10.0 ± 1.3 | 23.5 ± 3.6 |
| **BDL surgery 7 d** | -1.6 ± 1.4 | 10.4 ± 0.8 | 13.2 ± 0.5 |
| **ischemia-reperfusion** | -1.5 ± 0.4 | 7.7 ± 1.9 | 21.0 ± 7.6 |
| **APAP** | -0.4 ± 1.0 | 21.7 ± 19.8 | 13.6 ± 10.3 |
| **PCI** | -2.4 ± 0.7 | 0.9 ± 0.1 | 4.3 ± 0.4 |

**Supplementary Table S3:** Summary of disease parameters from human liver sections. CI: confidence interval; SD: standard deviation. * p<0.05 vs. reference group. For mean age, bilirubin, aspartate aminotransferase (ASAT), alanine aminotransferase (ALAT), C-reactive protein (CRP) the Mann-Whitney test was applied (data are not normally distributed). For albumin the unpaired t-test was performed (normality and equal distribution present).

| | **reference group**<br>(bilirubin $< 21 \mu\text{mol L}^{-1}$) | **cholestasis group**<br>(bilirubin $\geq 21 \mu\text{mol L}^{-1}$) |
| --- | --- | --- |
| **mean age ± CI** | 59 ± 4.9 years | 59 ± 10.1 years |
| **gender ratio** | 18 female, 16 male | 1 female, 7 male |
| **bilirubin ± SD** | 11.8 ± 4.1 $\mu\text{mol L}^{-1}$ | 64.7 ± 50.1 $\mu\text{mol L}^{-1}$ * |
| **ASAT ± SD** | 0.4 ± 0.2 $\mu\text{mol L}^{-1}$ | 0.8 ± 0.7 $\mu\text{mol L}^{-1}$ |
| **ALAT ± SD** | 0.5 ± 0.3 $\mu\text{mol L}^{-1}$ | 1.5 ± 2.3 $\mu\text{mol L}^{-1}$ * |
| **albumin ± SD** | 36.1 ± 5.6 g L<sup>-1</sup> | 28.6 ± 7.1 g L<sup>-1</sup> * |
| **C-reactive protein ± SD** | 37.7 ± 65.6 mg L<sup>-1</sup> | 69.8 ± 115.5 mg L<sup>-1</sup> |
| **disease entities included** | metastatic sigma carcinoma,<br>klatskin carcinoma,<br>adenocarcinoma (rectum,<br>ovary, colon, pancreas,<br>jejunum, stomage), metastatic<br>insulinoma, metastatic colon<br>carcinoma, breast carcinoma,<br>metastatic rectal carcinoma,<br>adenoma, liver hematoma,<br>hepatocellular carcinoma,<br>squamous cell carcinoma<br>cervix | metastatic colon carcinoma,<br>metastatic rectal carcinoma,<br>gall bladder carcinoma,<br>klatskin carcinoma,<br>neuroendocrine carcinoma,<br>adenocarcinom (pancreas),<br>metastatic cecum carcinoma |

